# Protein sequence landscapes are not so simple: on reference-free versus reference-based inference

**DOI:** 10.1101/2024.01.29.577800

**Authors:** Thomas Dupic, Angela M. Phillips, Michael M. Desai

## Abstract

In a recent preprint, Park, Metzger, and Thornton reanalyze 20 empirical protein sequence-function landscapes using a “reference-free analysis” (RFA) method they recently developed. They argue that these empirical landscapes are simpler and less epistatic than earlier work suggested, and attribute the difference to limitations of the methods used in the original analyses of these landscapes, which they claim are more sensitive to measurement noise, missing data, and other artifacts. Here, we show that these claims are incorrect. Instead, we find that the RFA method introduced by Park et al. is exactly equivalent to the reference-based least-squares methods used in the original analysis of many of these empirical landscapes (and also equivalent to a Hadamard-based approach they implement). Because the reanalyzed and original landscapes are in fact identical, the different conclusions drawn by Park et al. instead reflect different interpretations of the parameters describing the inferred landscapes; we argue that these do not support the conclusion that epistasis plays only a small role in protein sequence-function landscapes.

---

**I**n a recent manuscript (1), Park, Metzger, and Thornton apply methods they recently developed (2) to reanalyze 20 empirical protein sequence-function landscapes originally characterized by us (3–6) and by others (7–15). They argue that these protein sequence-function landscapes are relatively simple, with higher-order epistasis playing a less important role than earlier work suggested. Here, we show that the conclusions Park et al. draw from their reanalysis result from a misinterpretation of the relationship between different methods of characterizing protein sequence-function landscapes.

Park et al. highlight three key differences between their analysis and that used in the original characterizations of the 20 empirical landscapes they consider: (1) They claim that their “Reference-free analysis” (RFA) method is more accurate and more robust to measurement error and partial sampling than earlier approaches. (2) They allow for effects of global “nonspecific epistasis,” in which the measured protein function is a nonlinear function of some underlying phenotype. They introduce a simple form for this global nonlinearity, and claim that with this assumption the underlying phenotype has a simpler and less epistatic architecture. (3) Where appropriate they use a model with 20 possible alleles (interpreted as amino acid states) per locus, rather than a standard biallelic model.

Here we show that point (1) above is incorrect. As a basis for comparison with their RFA method, Park et al. implement a reference-based model that estimates coefficients without any averaging. This model, as they claim, is therefore highly sensitive to noise. However, this is not the approach used in most of the original analyses of the 20 empirical landscapes they consider. Instead, the reference-based framework actually used in the original characterization of most of these landscapes is implemented using linear regression. Here we show that these models are exactly equivalent to the RFA model implemented by Park et al. Thus the original analysis is exactly as robust to measurement noise and missing data as their reanalysis, and leads to identical results. This exact relationship is not a novel result, and is well established in the quantitative genetics literature (see *e*.*g*. (16)). Park et al. also claim their RFA framework is superior to a Hadamard-based approach used in the original analyses of a few of the empirical landscapes. However, the RFA and Hadamard-based models are also exactly equivalent, with coefficients that are identical up to a simple multiplicative factor which depends on the precise convention chosen for the Hadamard model (though Park et al. are correct to note that the Hadamard approach by construction cannot account for missing data (16)). We note that in some other contexts (*e*.*g*., when addition of additive or epistatic coefficients to a model is constrained by a regularization term), these various approaches can yield different results. In these cases, the relative performance of the different methods depends on the specific context, and no method can be considered better in an absolute sense.

We next turn to point (2) above. We argue that global epistasis can provide a simpler representation of protein sequence-function landscapes in some cases, particularly where there is a natural biochemical or biophysical reason to believe that protein function depends nonlinearly on some underlying additive phenotype. For example, in some cases we might expect mutations to affect free energy of folding or binding, which in turn are related nonlinearly to protein functions defined in terms of fraction of protein folded or bound. However, in many of the 20 empirical landscapes reanalyzed by Park et al., the measured phenotype is itself a free energy, and there is no biochemical basis for assuming that this is related nonlinearly to some other underlying additive phenotype. In such cases, we argue that introducing global epistatic functions can instead distort biological interpretations of the data.

We do not directly address point (3) in this manuscript. The use of 20-allele models is indeed appropriate when considering landscapes involving many or all possible amino acid variants at a given protein residue. However, in 12 of the 20 empirical protein landscapes reanalyzed by Park et al., there are only two possible alleles considered at each site. Thus for these landscapes, this aspect of the method is not relevant.

**Table 1.**
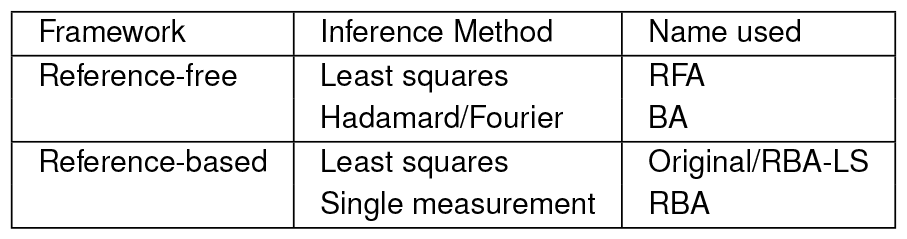
Overview of the epistasis inference methods considered.

| Framework | Inference Method | Name used |
| --- | --- | --- |
| Reference-free | Least squares | RFA |
|  | Hadamard/Fourier | BA |
| Reference-based | Least squares | Original/RBA-LS |
|  | Single measurement | RBA |

## Reference-based and reference-free epistasis frameworks are exactly equivalent

Consider a protein sequence-function landscape defined by all possible genotypes at some set of *L* loci. We consider here the biallelic case, so there are 2^*L*^ possible genotypes, each of which corresponds to some functional value *y* (though we note that Park et al. also consider a more general 20-allele model, which introduces additional complexities which we do not address here). In what Park et al. refer to as a “reference-free” framework, we write the landscape as

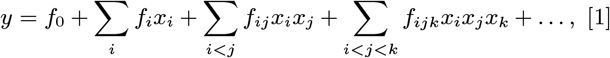

where *x*_*i*_ ∈ {+1,−1} is the genotype at locus *i*. Note that because there are as many coefficients *f* as there are genotypes, we can write an arbitrary sequence-function landscape in this form (though we may wish to truncate the landscape at some order *K*, in which case we refer to it as a linear epistasis model at order *K*). The choice of *x*_*i*_ ∈ {+1,−1} in Eq. (1) is what defines the reference-free framework (though as noted above this is specific to the biallelic case). It is simple to show that *f*_0_ is the average functional value, that *f*_*i*_ is half the effect of changing the allele at locus *i* (from−1 to +1), averaged over backgrounds at all other loci, that *f*_*ij*_ is half the average effect of changing the alleles at loci *i* and *j* from an asymmetric to a symmetric configuration, and so on. In this sense, the *f* terms can be thought of as reference-free, because there is no reference to any wild-type sequence and all the terms are defined instead based on average effects across all possible states at other loci. We often refer to the *f*_*i*_ as additive terms, the *f*_*ij*_ as pairwise epistatic terms, and so on, but it is important to remember their precise meaning as defined in Eq. (1).

Park et al. contrast reference-free frameworks with reference-based frameworks. In the reference-based framework, we instead write the landscape defined above as.

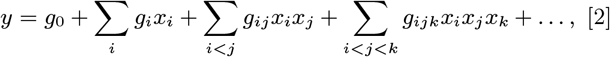

where now *x*_*i*_ ∈ {0, 1} As above, we can truncate this at order *K* to create a linear epistasis model at that order, or consider the full model with *K* = *L*. Due to this alternative choice of how the genotype is represented, the genotype with *x* = 0 at all loci now has a special status as the “wild-type” sequence, and at each locus we can refer to the “wild-type” allele as *x*_*i*_ = 0 and the “mutant” allele as *x*_*i*_ = 1. We can easily see from Eq. (2) that *g*_0_ is the functional value of the wild-type sequence, that *g*_*i*_ is the effect of changing locus *i* from the wild-type to the mutant allele on a background that is wild-type at all other loci, that *g*_*ij*_ is the effect of having the mutant allele at both loci *i* and *j* above and beyond the sum of the additive effects of mutant alleles at these loci (on a background that is wild-type at all other loci), and so on. We often refer to the *g*_*i*_ as additive terms, the *g*_*ij*_ as pairwise epistatic terms, and so on, but as above it is important to remember their precise meaning, in this case as defined in Eq. (2).

Because of the different representations of the landscapes in the reference-free versus reference-based frameworks, the *f* and *g* terms will not be the same. Thus our interpretation of what are called additive versus epistatic effects may be different. However we emphasize that this is the exact same landscape: exactly the same genotypes correspond to exactly the same functional values. As a simple example, consider a two-locus landscape with four possible genotypes: AB, aB, Ab, and ab. Imagine the functional values of each of these four genotypes are *y*_*AB*_ = 0, *y*_*aB*_ = 0, *y*_*Ab*_ = 0, and *y*_*ab*_ = 1. In the statistical framework we have *f*_0_ = 1*/*4, *f*_1_ = 1*/*4, *f*_2_ = 1*/*4, and *f*_12_ = 1*/*4, while in the biochemical framework we have *g*_0_ = 0, *g*_1_ = 0, *g*_2_ = 0, and *g*_12_ = 1. Thus in the reference-free framework we interpret this landscape as having both additive and pairwise epistatic terms, while in the reference-based framework we interpret it as having no additive term and a stronger pairwise epistatic term. However, the landscape is identical in both frameworks, and the difference is merely a matter of how the coefficients are interpreted. This is true regardless of what order the model is truncated at: any sequence-function landscape that can be represented at order *K* in the reference-free framework is exactly equivalent to a corresponding landscape in the reference-based framework at that same order *K*.

Park et al. argue that, although the true landscape may be identical in both the reference-free and reference-based frameworks, inferring its structure in the reference-free framework using their RFA approach is more accurate and more robust to measurement noise and partial sampling. They are indeed correct that RFA as they implement it is more robust than the reference-based inference method that they implement (which they term “RBA”). This is because their RBA method computes coefficients in the reference-based framework based on differences between specific reference genotypes (*i*.*e*., first-order terms are defined based on the differences in single-mutant sequences relative to the wild-type, second-order terms based on deviations from additivity in double-mutants, and so on). This simple method is extremely sensitive to measurement noise, as each coefficient is inferred based on very few phenotypic measurements. It is also very sensitive to missing data — for example, if a double mutant is absent, all higher order effects involving the corresponding two mutations cannot be estimated. Because of these weaknesses, this method cannot be recommended and is seldom used to study epistasis in empirical landscapes.

However, this poor performance is due to the limitations of the RBA inference method, not the reference-based framework itself. Instead of using RBA, most of the original analyses of the 20 empirical landscapes considered by Park et al. instead used least squares regression to infer coefficients within the reference-based framework (*i*.*e*., truncating the model to some order *K* < *L* and fitting the coefficients to minimize least-squared error in the observed phenotypes). We refer to this inference approach as RBA-LS. We show here that RBA-LS is exactly equivalent to RFA at any order *K*. Thus these original analyses are exactly as accurate and robust to measurement noise and partial sampling as the reanalysis of Park et al. The key point is that although the reference-based framework does make reference to a wild-type sequence, inference of the coefficients using least squares regression effectively averages each coefficient over many sequence backgrounds, and hence is not overly sensitive to measurement errors in any single sequence or set of sequences.

To see this, we represent the genotype at *L* biallelic loci as the vector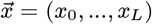 with *x*_*i*_ ∈ {0, 1}, and write an arbitrary protein structure-function landscape involving these loci as:

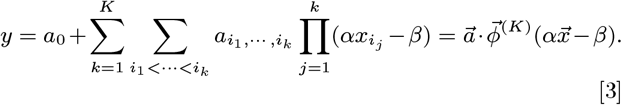

This landscape is linear in the coefficients *a* and has 2^*K*^ coefficients at order *K*. A complete model, where *K* = *L*, would contain as many coefficients as there are available genotypes (2^*L*^). The reference-free framework considered by Park et al. corresponds to *α* = 2 and *β* = 1, while the reference-based framework corresponds to *α* = 1 and *β* = 0.

Given a dataset of phenotypes 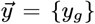 and associated genotypes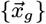, these models can be inferred by minimizing the least squares error. Because the models are linear, the solution can be computed explicitly as a function of *ϕ*^(*K*)^. We have

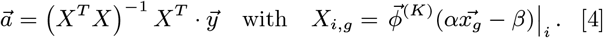

With this formalism, it is straightforward to see that the coefficients generated by one set of (*α, β*) parameters are linearly related to the coefficients generated by a different set of parameters: both models will give the same predictions and have exactly the same goodness-of-fit. This is true regardless of what order *K* the models are truncated at, and remains true regardless of the scale of measurement errors or missing data.

We emphasize that this equivalence does not only hold at the level of model performance or of predicted phenotypes: all of the coefficients inferred in either framework are also exactly interchangeable. That is, we can infer the reference-free coefficients by first using RBA-LS to infer the reference-based ones and then applying the linear transformation described above. The resulting reference-free coefficients are identical to those inferred by RFA directly. The converse is also true: RFA infers the same reference-based coefficients (after applying the linear transformation) as RBA-LS.

To illustrate this equivalence between the inferences in the reference-free and reference-based frameworks, we simulated a biallelic landscape with *L* = 12 sites with random (normally distributed) epistatic coefficients up to order 3 (see Methods for details). We added varied degrees of simulated experimental noise to this data. We then used the reference-free least squares method (*i*.*e*. RFA as defined by Park et al.) and the reference-based least squares method (as implemented in our original analyses of several of these empirical landscapes) to infer the landscape. As expected, we find that the landscapes inferred by the two methods are identical (Fig 1A), and explain identical amounts of phenotypic variance at each order (Fig 1E). This remains true regardless of the extent of simulated noise (Fig 1G). Provided that experimental noise is not too extensive, these inferences accurately recapitulate the true underlying landscape (Fig 1D); because both models lead to identical landscapes this goodness of fit is also identical. We also explored the effect of missing data by randomly removing some fraction of the simulated measurements; as expected, the reference-free and reference-based least squares inferences remain identical (Fig 1H). We also inferred the landscape using the subtraction-based (RBA) method implemented by Park et al, and find as expected that this approach is not equivalent to the other two methods (Fig 1C), explains much less of the phenotypic variance (Fig 1E), and is substantially more sensitive to experimental noise (Fig 1G). We implemented a standard cross-validation approach to find the optimal order epistatic model; both the reference-free and reference-based least squares approaches correctly select the simulated third order model (Fig 1F).

**Fig. 1.**
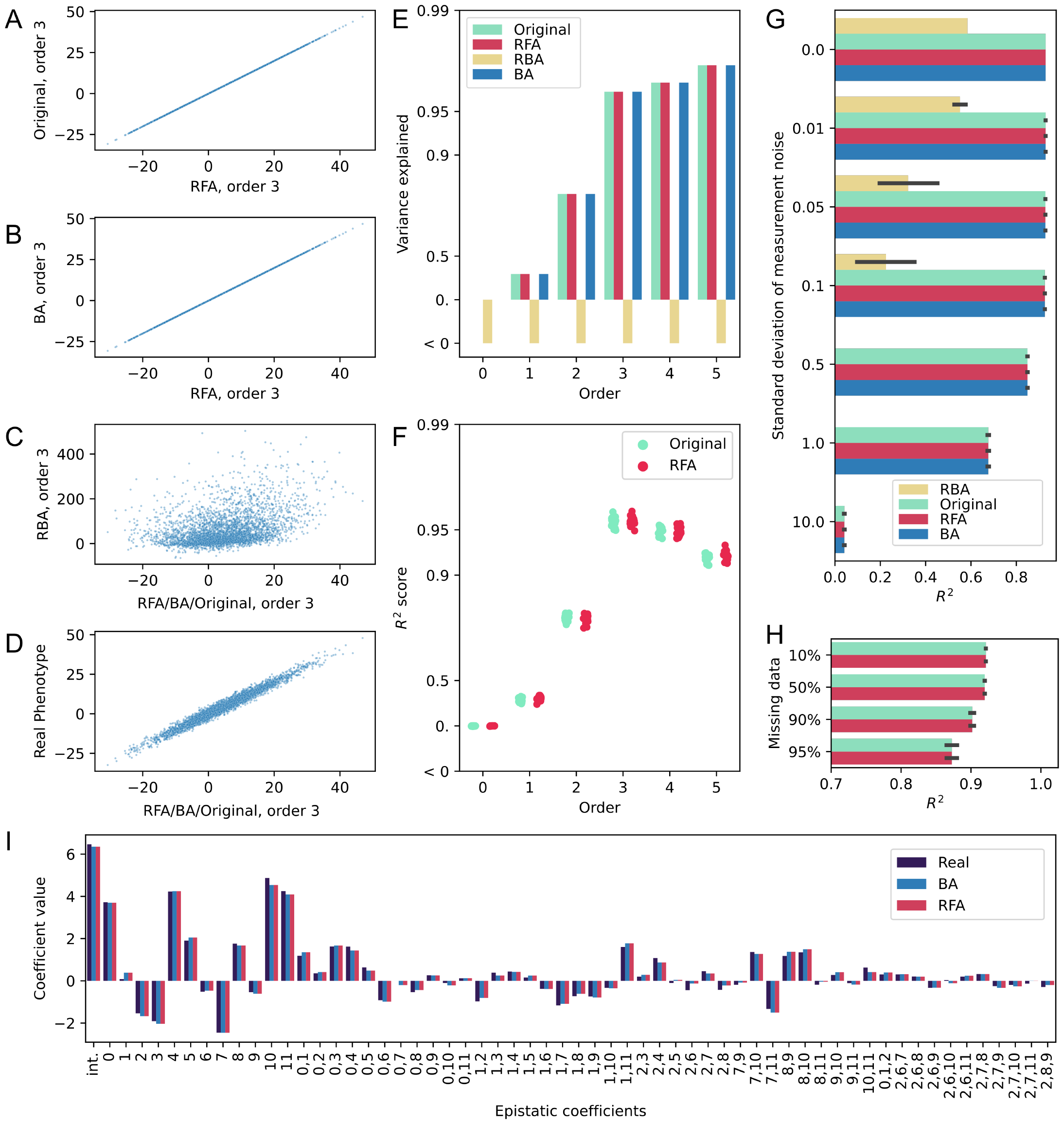
Performance of the different models in a simulated 12-site (*L* = 12 landscape; max epistatic order 3) that includes simulated experimental noise (see Methods for details). (**A-D**) Comparison of phenotype predictions between least squares reference-free (RFA) model and least squares reference-based model used in the original analysis of this data (**A**), Hadamard reference-free (BA) model (**B**), reference-based model without averaging (RBA) (**C**) and the actual phenotypes (**D**). Note that the RFA method is exactly equivalent to the least squares reference-based and BA approaches. (**E**) Variance explained by the different models at all orders. Note the exact correspondence between RFA, original, and BA approaches. (**F**) Cross-validation for the original and RFA approaches (the two least squares methods compatible with missing data), trained on 80% of the data with *R*^2^ evaluated on the remaining 20%. (**G**) The effect of simulated measurement noise on the *R*^2^ of the models inferred using the different approaches. Note that the original least squares reference-based method performs identically to RFA and the BA approach regardless of the amount of experimental error. (**H**) Effect of missing data on the *R*^2^ of the inferred models. Note that the original and RFA approaches (the two methods compatible with missing data) perform identically regardless of the fraction of data missing. (**I**) Comparison between a subset of the epistatic coefficients obtained with the two reference-free methods (RFA, BA) and the real coefficients. Note the exact correspondence between RFA and BA methods.

We next repeated this comparison for our CR9114-H1 data, as one example from the 20 landscapes reanalyzed by Park et al. Again, we find that the reference-free and reference-based least squares models infer identical landscapes (Fig 2A), and explain identical amounts of phenotypic variance at each order (Fig 2E). This inferred landscape provides reasonably accurate predictions of the observed phenotypes (Fig 2D). In contrast, the simple reference-based method (RBA) leads to very different inferences (Fig 2C), which are much less accurate and perform very poorly in explaining phenotypic variance (Fig 2E). A standard cross-validation approach with the least squares methods finds that the fifth-order model is optimal, as we found in our original analysis (Fig 2F).

**Fig. 2.**
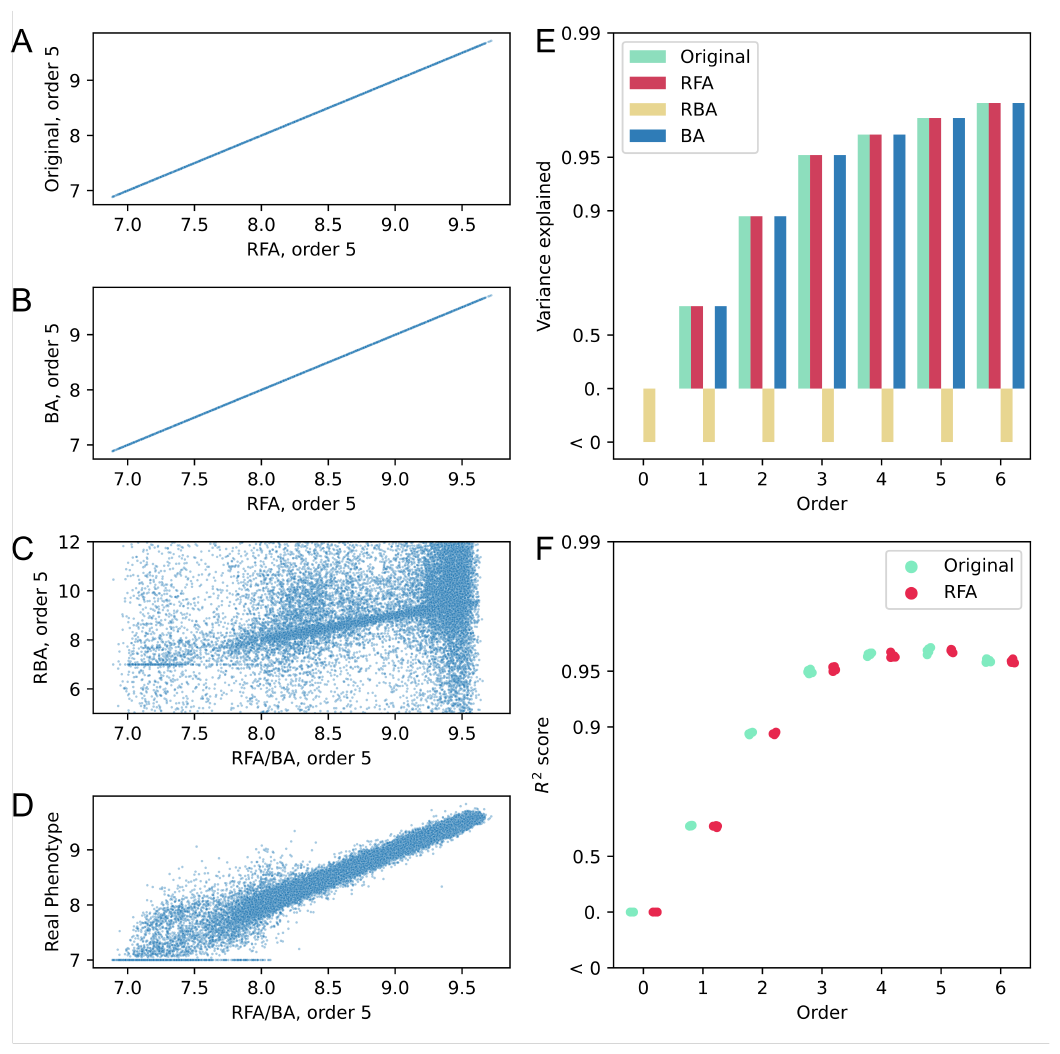
Performance of the different models on the empirical CR9114-H1 dataset. (**A-D**) Comparison of phenotype predictions between least squares reference-free (RFA) model and least squares reference-based model used in the original analysis of this data (**A**), Hadamard reference-free (BA) model (**B**), reference-based model without averaging (RBA) (**C**) and the actual phenotypes (**D**). Note that the RFA method is exactly equivalent to the least squares reference-based and BA approaches. (**E**) Variance explained by the different models at all orders. Note the exact correspondence between RFA, original, and BA approaches. (**F**) Cross-validation for the original and RFA approaches (the two least-squared methods compatible with missing data), trained on 80% of the data with *R*^2^ evaluated on the remaining 20%.

## The reference-free least squares and Hadamard models are exactly equivalent

Park et al. also compared their reference-free least squares model (RFA) to a Hadamard model (this has also been called a Fourier or Hadamard-Walsh model (16)), and more particularly to a specific implementation of this which they term a “background-averaged” (BA) model. In this comparison, Park et al. claim that, for most metrics, the reference-free least squares model outperforms the Hadamard model. Contrary to this assertion, we show here that the Hadamard-Walsh transformation and the least squares approach are equivalent. This equivalence has been described in previous work (16), with the apparent discrepancy found by Park et al. arising from a failure to account for a scaling difference in how the epistatic coefficients are defined in the two models (16).

To clarify this difference in scaling factor, we rederive the Hadamard matrix here. We write the phenotype *y* associated with genotype (*k*_1_,, *k*_*L*_) (with *k*_*i*_ ∈ {0, 1}) as 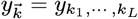.We call 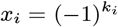 the genotype in the reference-free (*x*_*i*_ ∈ {+1*/* −1}) framework. The multi-dimensional discrete Fourier transform of this phenotype can then be expressed as

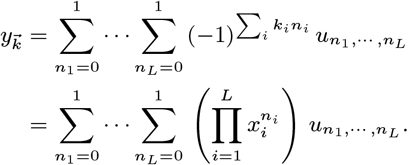

We then define 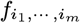 (with *i*_1_ *<* … *<i*_*m*_) such that 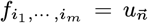 with *n*_*i*_ = 1 if *i* ∈ {*i*_1_, ⃛ *i*_*m*_} else 0. For example, with *L* = 3, *f*_2_ = *u*_010_ and *f*_12_ = *u*_110_. Rewriting the above equation, we find at maximum order *K* = *L*:

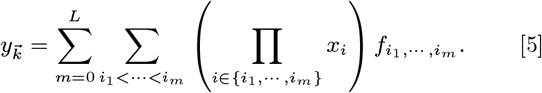

We see from this result that for the complete model (*K* = *L*), the coefficients *f* of the reference-free framework are identical to the Fourier coefficients *u*, but with a different indexing system. The advantage of the Hadamard method is that the coefficients *u* are easier to manipulate algebraically and can be obtained by computing the inverse Fourier transform on the phenotypes:

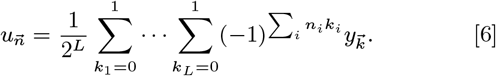

This is known as the Walsh-Hadamard transformation, and can be written in matrix form as 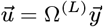 where the matrix Ω^(*L*)^ is usually given as a recursion formula in the number of sites *L*, which we can find by fixing the value of the first site:

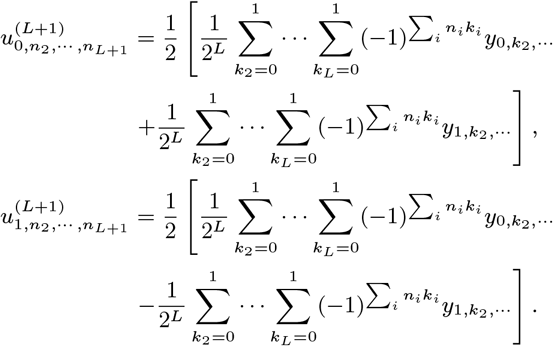

This leads to the recursion relation:

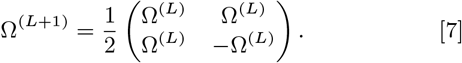

Up to a factor of 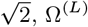 is the standard Hadamard matrix. However, this formula differs from the one used by (1), 16) and (13). Instead, the definition quoted in these papers adds a factor of (−2)^K^ to the coefficients *f* at order *K*. This discrepancy was not compensated for by Park et al. when comparing the two models. Because their Hadamard coefficients are a factor of 2^K^ larger than the RFA coefficients, the error on these coefficients is also a factor of 2^K^ larger.

Correcting for this factor, we find that the RFA and Hadamard methods are identical, resulting in the exactly the same landscape and parameterized by exactly the same coefficients (appropriately re-indexed and scaled). This is true regardless of what order the models are truncated at. Specifically, the Walsh-Hadamard transformation makes it possible to directly compute the coefficients of the reference-free framework for the full order model (with *K* = *L*). Because (as noted by Park et al.) the coefficients in the reference-free framework remain constant regardless of model order, the coefficients of a lower order reference-free least squares model (i.e. with *K < L*) also exactly match a corresponding subset of the coefficients inferred using this reference-free Hadamard approach. This exact correspondence can be verified numerically (see Fig. 1B, 1I, and 2B). The Hadamard method can therefore be preferable to the least squares method, as it is often faster and yields the exact same coefficients. This remains true in the presence of any amount of noise in the experimental measurements.

However, the main drawback of the Hadamard approach is missing data. The method will only perform correctly at order *K* if the data is complete up to order *K* (16). Park et al. attempted to address this by applying linear regression with elastic net regularization to generate what they call their “background-averaged” (BA) method, and make a number of comparisons between RFA and BA. As they noted, the two methods perform similarly in the two-sites scenario, which is unsurprising as they are both essentially the same linear regression, fitted with either least squares (for RFA) or elastic net (for their “BA”). It should also be noted that in presence of missing data, the reference-free least squares approach loses its distinctive statistical properties. Notably, the average of the phenotypes no longer serves as the intercept, and the model coefficients become dependent on the maximal order of the inferred model. This is because missing data breaks the connection between the least squares approach and the Hadamard/Fourier transform (16). It is also important to note that the equivalence between these two approaches no longer applies when least squares estimation is modified to include regularization terms (*e*.*g*., lasso or ridge regression; see below for further discussion).

## On interpreting the inferred landscapes

We have seen above that reference-free least squares landscape inferred by Park et al. in their reanalysis of our CR9114-H1 data (and many of the other 19 empirical data sets they consider) is identical to the reference-based least squares landscape we originally inferred (up to considerations involving “global” epistasis, which we return to below). The different conclusions Park et al. draw about the extent and importance of epistasis therefore reflect the different interpretations of the coefficients in the two frameworks, rather than any true difference in the inferred landscapes.

What are the key differences in interpretation between the coefficients of the reference-free versus the reference-based framework? In reference-free least squares model, the coefficients have the desirable property of being summary statistics for the phenotype distribution. For example, *f*_0_ represents the mean phenotype value, and *f*_*i*_ is half the average effect of the *i*th mutation. Because of this statistical interpretation, and as Park et al. have noted, these coefficients do not depend on the order *K* at which the inferred model is truncated. However, the tradeoff for this desirable property is that, as ensemble values, the reference-free least squares coefficients are sensitive to the set of loci considered in the model. For example, if we decide to fix one of the mutations, or to add an additional locus to the data set, the reference-free least squares coefficients will all change. In that sense, calling these coefficients reference-free is misleading, because they depend on which mutations are considered in the dataset.

By contrast, the reference-based least squares coefficients capture interactions between mutations more directly. For example, in our CR9114 and CR6261 datasets, we directly measure binding free energy Δ*G* to hemagglutinin of each antibody sequence variant. Because we expect these energies to be additive, a natural null expectation is that a given mutation *i* makes some change ΔΔ*G*_*i*_, potentially with some interactions with other mutations. While this model is relatively simple (*e*.*g*., it does not take into account large structural changes), the coefficients of the reference-based least squares epistatic model will correspond to these changes in binding free energy. However, because the coefficients are not independent of the order of the model, this interpretation is only valid if the model already captures most of the relevant epistatic interactions.

These considerations highlight the fact that the reference-free and reference-based epistasis frameworks serve distinct purposes. Reference-free epistasis is often employed in genomics for phenotypes that are not directly biochemical (*e*.*g*., fitness and other complex traits) (16). In this case, any biochemical meaning is usually lost. It is particularly relevant in considering evolution in outcrossing populations, where the model coefficients are then directly related to average selection pressures. On the other hand, when looking at protein phenotypic properties, reference-based epistasis can be more relevant. It is also more useful in analyzing mutational trajectories, where model coefficients are directly related to selective effects of individual substitutions. However, both models will result in identical fit, and the coefficients of one can be transformed easily into the coefficients of the other.

In both the reference-free and reference-based frameworks, the number of coefficients increases exponentially with the maximum order of the model, *K*. With more coefficients and the same number of phenotypes, the impact of phenotype measurement errors on the inferred coefficients also increases exponentially. Specifically, and in both frameworks, the error in the highest-order coefficients grows as 2^*K/*2^, where *K* is the order of the model. Where the two frameworks differ is that in reference-free least squares epistasis, low-order terms remain stable regardless of the model order, meaning their associated errors do not increase. By contrast, in reference-based least squares epistasis, all the coefficient errors generally increase with model order, at a speed that depends on the epistatic structure. However this does not affect the error on the inferred phenotypes (see Fig 1E, Fig 2E) – the two models are still exactly identical, the error is simply distributed differently across coefficients. Because both sets of coefficients can be interconverted without any loss of precision, the choice of representing in one framework rather than the other cannot be made based on expected error, and instead should be guided by the biological question at hand.

A key overall conclusion of Park et al.’s reanalysis is that protein sequence-function landscapes are simpler and less epistatic than earlier work suggested. This conclusion rests primarily on their finding that variance explained (or equivalently *R*^2^ of the predicted versus measured phenotypes) increases only slightly with increasing model order. However, as we have seen, the exact same conclusion can be drawn from our original reference-based least squares analysis of the CR9114-H1 dataset (Fig. 2F), along with many of the 19 other empirical landscapes. Why then did earlier work argue that high-order epistasis is important? The basic reason is that variance explained is not the only measure of importance: high-order epistasis can be widespread and significant while still not substantially increasing variance explained. This is particularly clear from our simulated data. In these simulations, the third-order terms have the same total magnitude as the pairwise and additive terms. They are therefore widespread and significant, and and will certainly have a dramatic impact on the accessible evolutionary trajectories across this landscape. Yet adding these terms only slightly increases variance explained and cross-validated *R*^2^ (Fig. 2E,F note the logarithmic scale).

The relationship between variance explained, crossvalidated *R*^2^, and model order for the CR9114-H1 dataset (and many of the other 19 empirical landscapes) has a similar form to our simulated data. Thus it entirely plausible that high-order terms are widespread and significant, as claimed in the original analysis of this data, even though their contribution to variance explained is limited. This is supported both by the cross-validation and by our estimates of the measurement errors in high-order coefficients in our original analysis. For example, if *σ* is the standard error in our phenotype measurements, the error in the coefficients in our original reference-based least squares inference of the CR9114-H1 landscape at order 5 is 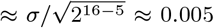. This is negligible compared to the value of the epistatic coefficients we deemed significant (some order 5 coefficients reach 0.5; see Fig 3F in (3)). Thus this epistasis is well supported by the data and is potentially both biochemically and evolutionarily significant, despite not contributing substantially to overall phenotypic variance explained.

**Fig. 3.**
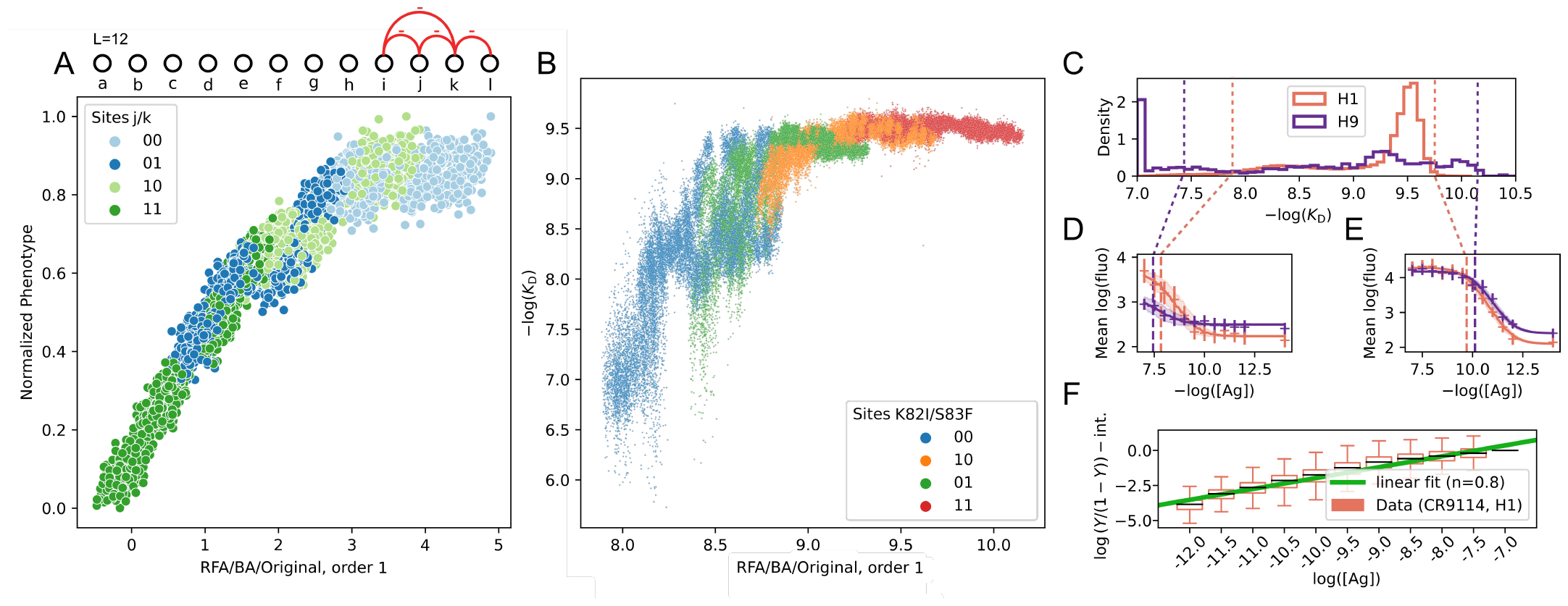
(**A**) Comparison of a first-order model with normalized phenotype in a simulated 12-site, second-order landscape. The first-order coefficients are randomly generated and only four second-order biochemical coefficients are non-zero (see graph above). (**B**) Measured binding affinity versus predictions of the first-order epistasis model for our CR9114-H1 empirical landscape, with genotype at two specific loci highlighted. (**C**) Binding affinity distribution for CR9114-H1 data (orange) and CR6261-H9 data (purple). (**D**) Two example titration curves for H1 (orange) and H9 (purple) with high *K*_D_ values. Cross markers show inferred fluorescence levels, while the fitted curve is represented by the line. A vertical dashed line indicates the *K*_D_ value derived from the fit. (**E**) Similar representation for examples with low *K*_D_ values. (**F**) Hill plot for CR9114-H1, including Hill coefficient fit result (*n* = 0.8).

Finally, we emphasize that the equivalence between the reference-free and reference-based epistasis frameworks only holds if coefficients are inferred through least square minimization. This is what is used by Park et al. for their reference-free analyses (RFA), and what was used by us in the original analysis of our datasets. The reference-based approach that they compare their RFA model to infers coefficients recursively, beginning by subtracting the single reference phenotype from each individual measurement. The coefficients generated by this method indeed have more uncertainty, because they depend sensitively on measurement errors in one or a few reference strains (Fig1C,E, Fig2C,E). In most cases this approach cannot be recommended.

We also note that because the coefficients represent different quantities (despite the overall model being the same), the reference-free and reference-based frameworks will no longer lead to exactly identical models if there is some regularization imposed on the inference of the coefficients. For example, if one uses Lasso or elastic net regularization, rather than plain linear regression, both models can perform differently. However, in these cases the model that performs better will be the one for which the regularization is more appropriate (*i*.*e*., for any particular landscape the regularization penalty for the true model coefficients will be smaller in one or the other framework; that framework will perform better for that landscape). It is our view that when the measured phenotype reflects some underlying biochemical property (*e*.*g*., a free energy of binding or folding) it is more appropriate to regularize for sparsity in the reference-based framework, because we expect interactions between sites to only occasionally affect these properties. When the measured phenotype is some more complex phenotype, this may not be true. However, this is merely speculation, and much further empirical work will be needed to determine whether one approach is typically better or if the appropriate framework for regularization tends to vary depending on the specific context.

## Global versus idiosyncratic epistasis

Beyond the formalisms described above, an alternative approach to describing sequence-function landscapes is to view deviations from additivity as arising from a nonlinear transformation of an underlying, unobserved additive phenotype. In this approach, non-additive phenotypes can be transformed into their underlying additive counterparts by applying a simple nonlinear transformation, which represents the “nonspecific” or “global” epistasis present in the dataset (17, 18). In principle, this transformation can capture (1) the relationship between the measured phenotype and an underlying additive phenotype, (2) measurement noise or limits on the dynamic range, and (3) a combination of specific epistatic effects, like those described above. Following transformation, any remaining deviations from additivity are attributed to true specific epistatic effects.

This approach has been widely implemented by the protein evolution field, and these studies typically observe fewer significant epistatic coefficients following the non-linear transformation (19–21). Mathematically, this result is obvious: specific (“idiosyncratic”) epistatic effects can always be captured by a global transformation, and global epistasis can always be represented by specific epistatic coefficients. The question then becomes: which representation of the genotypephenotype architecture is more meaningful?

As an example of a system where idiosyncratic epistasis is the correct formalism, we simulated a twelve-site (*L* = 12) sequence-function landscape featuring only additive terms plus four pairwise epistatic coefficients (and no global nonspecific epistasis). We inferred a first-order landscape using the equivalent reference-based and reference-free least squares approaches. We then compare the predictions from this firstorder (additive-only) model to the simulated phenotypes. We find that a noticeable non-linearity emerges (Fig 3A). It would be natural to misconstrue this nonlinearity as a limitation in the dynamic range of the measurement and attempt to model it using a global nonlinear transformation. However, in this case applying the nonlinear transformation would obscure the genuine interactions and reduce the accuracy of the first-order linear coefficients.

This example illustrates why, in contrast to Park et al., we argue that whether a model of nonspecific epistasis is appropriate depends on the biological question. In cases where the measured phenotypes are on an arbitrary or relative scale (*e*.*g*., some measure of “fitness”), a nonlinear transformation may be appropriate to reveal an unmeasured additive phenotype (17). In contrast, in cases where the measured phenotypes are a thermodynamic state variable (*e*.*g*., free energy of binding), it is natural to expect that they behave additively without any such transformation (22). Applying a nonlinear transformation to this type of phenotype can still potentially reduce experimental noise or specific epistatic effects. However, rather than capture some unmeasured additive phenotype, this transformation may instead distort the phenotypic data and obscure its biological interpretation. Thus, there is an inherent trade-off to transforming additive phenotypic data: the ideal inference will reduce noise without removing biological features.

Even when the dynamic range of the experimental phenotype measurements is limited, a global epistasis inference may not be the best choice. Whenever possible, it is more appropriate to directly measure the experimental nonlinearities, and use this measured function. When this is not possible, and when the dynamic range of the experiment is well-defined, a censored regression model (23) (also known as a Tobit model) is a more appropriate way to capture information from phenotypes outside the dynamic range. This approach avoids the risk that fitting an arbitrary nonlinear function to the data would distort the information within the range. This approach is particularly useful when, for example, large parts of a protein binding affinity landscape consists of sequences that have no detectable binding; we used it for this reason in our analysis of a SARS-CoV-2 RBD landscape (6). Alternatively, if specific mutations cause genotypes to fall outside the experimental dynamic range, those mutations can be excluded from the epistatic inference, and can be interpreted as essential for having some measurable phenotype (as we did in our original analyses of the CR9114-H3, CR9114-FluB, and CH65-SI06 landscapes (3, 4)).

The trade-off between inference of global epistasis versus idiosyncratic effects will vary for each dataset and can be evaluated empirically, as we did in Phillips et al. 2021 ((3); see Appendix 2), by determining the appropriate nonlinear transformation and inferring specific epistatic effects on both transformed and untransformed data. For these datasets, we find that the phenotypes, which are free energies of binding, can be transformed onto an “additive” scale through a logistic function (similar to the function described by Park et al). Following logistic transformation, the optimal epistatic models are quite similar to those for the untransformed data, differing at most by one order. Because the transformed data did not produce substantially more concise models, we examined how the logistic transformation was altering the structure of the data. Across the datasets, there are many genotypes with exceedingly low or high affinity, and the logistic transformation distributes these genotypes to remove nonlinearity from the data. Though this transformation could be removing a technical artifact from the data (*e*.*g*., limited dynamic range), this is unlikely because affinities below the dynamic range correspond to non-specific binding (24), and the maximum phenotype is well within the dynamic range (Fig 3C,D,E) (3, 25). Furthermore, the aggregated Hill plots from our data (Fig 3F) validate that the Hill equation we employ to derive the dissociation constant is a good fit across the range of concentrations we investigate. Conversely, in several cases, the “pileup” of genotypes at low and high affinity corresponded to the presence of one or more specific mutations (Fig 3B). Thus, rather than reducing error, in this case we found that the global transformation distorted biological interpretations of the genotype-to-phenotype map.

## Discussion

We have shown here that least squares inference within the reference-free framework (Park et al.’s RFA model) and least squares inference within the reference-based framework (used in the original analysis of many of the 20 empirical protein sequence-function landscapes) lead to exactly equivalent sequence-function landscapes. We emphasize that this relationship has been demonstrated previously (see *e*.*g*. (16)); we simply rehash it here in the context of the recent work by Park et al. (1). As we discuss above, the choice between these models is primarily a question of how the coefficients are interpreted. Beyond this, the models are exactly equivalent: the inferred coefficients can be interconverted using a simple linear transformation, and they are identically susceptible to noise and missing data. Both models are far superior to the subtraction-based RBA method, which is very sensitive to experimental noise (and which was not used in the original analysis of most of the empirical datasets). We also show that Park et al.’s RFA model is identical to the “background averaged” Hadamard approach, with the deviations they report resulting from a difference in conventions.

Although the RBA-LS and RFA approaches are exactly equivalent inference methods, and lead to identical sequence-function landscapes, it is important to note that the reference-based and reference-free coefficients have different interpretations and are inferred with different degrees of precision. As Park et al. show, the reference-free coefficients can typically be estimated with higher accuracy than the reference-based coefficients, at least for the classes of sequence-function landscapes they focus on in their simulations. It may seem paradoxical that inference methods written in the two frameworks can be equivalent, and yet one set of coefficients is inferred (by either method) less accurately than the other. However, this is actually a common phenomenon in statistical inference. Consider for example a simple toy example: imagine we make many measurements of two independent quantities, *X* and *Y*. We then use least-squares inference to infer *X* and *Y* (call this method XY-LS), or alternatively to to infer Z = X+Y and W = X-Y (call this method ZW-LS). Under certain standard assumptions about the distributions of errors in our measurements, our estimates of *X* and *Y* will typically be more accurate than our estimates of *Z* and *W*. However, the two sets of coefficients are of course exactly equivalent; we can interconvert them easily and it is clear that XY-LS and ZW-LS are equivalent methods. The estimates of *Z* and *W* are less accurate because the errors in these quantities are anticorrelated, so when we convert them to *X* and *Y* we reduce the error.

As in this toy example, reference-based and reference-free coefficients are related by linear transformations, so it is not surprising that some sums or differences of quantities have higher or lower relative error. In fact, inferences of the reference-based coefficients (either by RBA-LS or RFA followed by conversion) are often not only less accurate, but also biased (i.e. they will not converge to the true values as the amount of data used for inference increases). It is tempting to argue that it is therefore somehow “better” to characterize protein sequence-function landscapes in the reference-free framework. However, the reference-based coefficients are physical properties of the protein (e.g. in our CR9114 datasets the first-order terms represent the effects of specific initial amino acid substitutions on antigen binding affinity). The fact that these physical properties can be estimated less accurately than other properties of the protein (e.g. the reference-free coefficients) is interesting, but does not imply that the reference-free framework provides the best way of interpreting the protein architecture. For example, the epistasis relevant for evolution from a germline to somatic antibody is directly related to the reference-based and not the reference-free coefficients, so the simplicity or complexity of the landscape is best assessed in those terms. By contrast, the epistasis relevant for evolution involving recombining loci in an outcrossing population can be more directly related to the reference-free coefficients, so in these cases it is more appropriate to assess the simplicity of the landscape in those terms.

Prior to implementing either the reference-based or reference-free inference approach, it may be necessary to transform the measured phenotypes onto an additive scale, particularly if the phenotype is not expected to behave additively or if the measurements are noisy and/or are affected by limited experimental dynamic range. If such a transformation dramatically reduces the complexity of the resulting model, the resulting more concise model may have improved biological interpretability. However, apparent global effects can also be caused by a few specific idiosyncratic interactions, in which case misinterpreting them as a nonlinear transformation can obscure important effects. Because of these considerations, in determining whether a nonlinear transformation of the data is appropriate, we recommend (1) considering whether the data are expected to behave additively, (2) confirming that phenotypes fall within the assay dynamic range, (3) determining whether the nonlinearity is attributable to a specific epistatic interaction, and (4) evaluating whether the transformation produces a more concise model.

In our view, our results show that high-order epistasis is indeed critical to understanding many of the 20 landscapes reanalyzed by Park et al., contrary to their overall conclusion. In some cases, we concur that almost all phenotypic variance can indeed be explained with models that include only additive effects and pairwise epistasis. However, as explained above, this does not mean that high-order terms are not widespread or that they do not play a critical role in shaping the space of accessible evolutionary trajectories. In other cases, where a substantial fraction of sequences lead to nonfunctional proteins (with phenotypes below the experimental threshold of detection), we argue that this is often indicative of strong high-order epistasis rather than a limited dynamic range, contrary to the suggestion of Park et al. This is likely to be particularly critical for mutational landscapes relevant for the gain of a new function, such as the formation of protein-protein interfaces that involve contacts between multiple residues, with individual contacts insufficient for binding. In such cases, we expect several mutations to be individually neutral but in combination to lead to detectable binding. This is arguably a particularly important form of high-order epistasis, and can easily cause a large fraction of genotypes to have phenotypes at the lower threshold for detectability (as in our CR9114-FluB dataset). Obscuring this effect (*e*.*g*., by incorrectly inferring strong additive effects for each mutation that lead to unphysically weak binding affinities, and then applying a nonlinear transformation to map these to the lower limit of detection) masks a key feature of the landscape that is critical for understanding constraints on evolution.

## Methods

For more in-depth methods, refer to Phillips, Lawrence et al. Appendix 2 (3). Here, we focus on the key elements of the above analyses. For Fig2A-F, we use the CR9114-H1 dataset (3). Filtered *K*_D_ values can be found on the original github. The code used to generate the figures is here.

Both least square models are inferred by numerically solving the linear equations 1 or 2. We employ the ‘scipy.linalg.lstsq’ function (26), with the ‘gelsd’ solver. For cross-validation, we randomly choose 80% of all genotypes for training and assess the *R*^2^ score on the remaining genotypes. The model at order *K* was inferred by solving equation 2 using only the phenotypes with *K* or less than *K* mutations. The resulting system is fully determined with as many equations as unknowns. This formulation is equivalent to (and simpler to implement than) the one described in Park et al.

The generated model in Fig 1 features *L* = 12 sites. The intercept follows a normal distribution centered at 2 with a standard deviation of 1. Its *n*-th order coefficients (reference-based) are normally distributed with a mean of 0 and a standard deviation calculated as 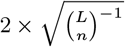 The scaling of epistatic coefficients is selected such that the sum of all coefficients at order *n* is independent of *n*. Phenotypes are generated from these coefficients, with a small added noise (normal distribution, standard deviation 0.05, except in Fig 1G,I). For Fig 1I, the noise level is increased to better show the difference between real and inferred values (standard deviation 10).

The model in Fig 3 has *L* = 12 sites with a randomly generated intercept (mean 2, scale 1) and first-order coefficients (mean 0, scale 0.2). Only four second-order coefficients are non-zero (i-j, j-k, k-l, i-k), each set to *−* 1.

The equation for the fitted curve in Fig 3D and Fig 3E is:

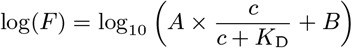

Here, *F* is the fluorescence, *c* is the concentration, and *A, B*, and *K*_D_ are the estimated coefficients. Mean receptor occupancy is given by 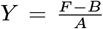 and the value plotted in Fig 3F is:

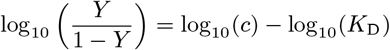

We adjust the intercept such that 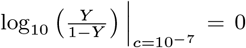 making the curves overlap regardless of *K*_D_ value. The slope of the curve gives the Hill coefficient (27).

## Acknowledgements

We thank Alief Moulana and other members of the Desai lab for helpful comments on the manuscript. TD acknowledges support from the Human Frontier Science Program Postdoctoral Fellowship, AMP acknowledges support from the Howard Hughes Medical Institute Hanna H Gray Postdoctoral Fellowship, and MMD acknowledges support from grant PHY-1914916 from the NSF and grant GM104239 from the NIH. Computational work was performed on the FASRC Cannon cluster supported by the FAS Division of Science Research Computing Group at Harvard University.

